# Investigation of contributions from cortical and subcortical brain structures for speech decoding

**DOI:** 10.1101/2023.11.12.566678

**Authors:** Hemmings Wu, Chengwei Cai, Wenjie Ming, Wangyu Chen, Zhoule Zhu, Chen Feng, Hongjie Jiang, Zhe Zheng, Mohamad Sawan, Ting Wang, Junming Zhu

**Author notes:** Corresponding authors: Hemmings Wu, Department of Neurosurgery, Zhejiang University School of Medicine Second, Affiliated Hospital, Hangzhou, Zhejiang, China, Ting Wang, School of Foreign Languages, Tongji University, Shanghai, China, Center for Speech and Language Processing, Tongji University, Shanghai, China, Junming Zhu, Department of Neurosurgery, Zhejiang University School of Medicine Second, Affiliated Hospital, Hangzhou, Zhejiang, China. These authors contributed equally to this work (Hemmings Wu, Chengwei Cai, Wenjie Ming, Wangyu Chen).

## Abstract

Language impairments often arise from severe neurological disorders, prompting the development of neural prosthetics based on electrophysiological signals for the restoration of comprehensible language information. Previous decoding efforts have focused mainly on signals from the cerebral cortex, neglecting the potential contributions of subcortical brain structures to speech decoding in brain-computer interfaces (BCIs). This study aims to explore the role of subcortical structures for speech decoding by utilizing stereotactic electroencephalography (sEEG). Two native Mandarin Chinese speakers, who underwent sEEG implantation for pharmaco-resistant epilepsy, participated in this study. sEEG contacts were primarily located in the superior temporal gyrus, middle temporal gyrus, inferior temporal gyrus, thalamus, hippocampus, insular gyrus, amygdala, and parahippocampal gyrus. The participants were asked to read Chinese text, which included 407 Chinese characters (covering all Chinese syllables), displayed on a screen after receiving prompts. 1-30, 30-70 and 70-150 Hz frequency band powers of sEEG signals were used as key features. A deep learning model based on long short-term memory (LSTM) was developed to evaluate the contribution of different brain structures during encoding of speech. Prediction of speech characteristics of consonants (articulatory place and manner) and tone within single words based on the selected features and electrode contact locations was made. Cortical signals were generally better at articulatory place prediction (86.5% accuracy, chance level = 12.5%), while cortical and subcortical signals predicted articulatory manner at similar level (51.5% vs 51.7% accuracy, respectively, chance level = 14.3%). Subcortical signals generated better prediction for tone (around 58.3% accuracy, chance level = 25%). Superior temporal gyrus remains highly relevant during speech decoding for both consonants and tone. Prediction reached the highest level when cortical and subcortical inputs were combined, especially for tone prediction. Our findings indicate that both cortical and subcortical structures can play crucial roles for speech decoding, each contributing to different aspects of speech.

## INTRODUCTION

Recently, there has been a significant interest in Brain-Computer Interfaces (BCIs) that can interpret speech from brain signals, potentially aiding those unable to speak^1, 2^. Understanding the mechanism of speech generation in the brain, including the sequence and location of involved brain regions, is crucial for developing a speech neuroprosthesis.

Current methods can decode text representations from neural signals during actual speech generation, spanning phonemes, words, full sentences, and even keywords. Many of these advancements utilize neural signals from cortical regions, including electrocorticography (ECoG) and Utah array, to record neural activity with high precision in time and space. While there are models explaining speech generation, the exact involvement of all brain regions remains unclear. Research now suggests that deeper brain areas like the hippocampus and thalamus play a role in both language comprehension and speech generation.

Stereotactic EEG (sEEG) is another commonly used surgical technique to record intracranial neurophysiological signals, where electrodes are implanted through small openings in the skull for treatment of refractory epilepsy. Unlike ECoG, which only records in cortical regions, sEEG is able to sample various regions, including subcortical brain structures, potentially benefiting BCI applications utilizing distant and deep brain areas. Here, we hypothesize that neural signals from subcortical brain regions can contribute to speech decoding. To validate our hypothesis that subcortical brain regions contribute to speech decoding, we asked participants to vocalize all possible pronunciation of characters in Mandarin Chinese while both their voices and sEEG data were recorded.

## MATERIALS AND METHODS

Two native Mandarin Chinese speaking patients with refractory epilepsy underwent sEEG surgeries. sEEG electrodes were implanted in cortical structures including superior temporal gyrus, middle temporal gyrus, and inferior temporal gyrus, and subcortical structures, including thalamus, hippocampus, insular gyrus, parahippocampal gyrus, and amygdala (Figure 1). The positions of the electrodes were confirmed manually by merging postoperative CT with preoperative MR. As the majority of the electrodes were located in the right hemisphere, electrodes in the left hemisphere were not included in this study. This clinical trial was approved by the Ethics Committee of the Zhejiang University School of Medicine Second Affiliated Hospital (protocol number: I2022145).

**Figure 1.**
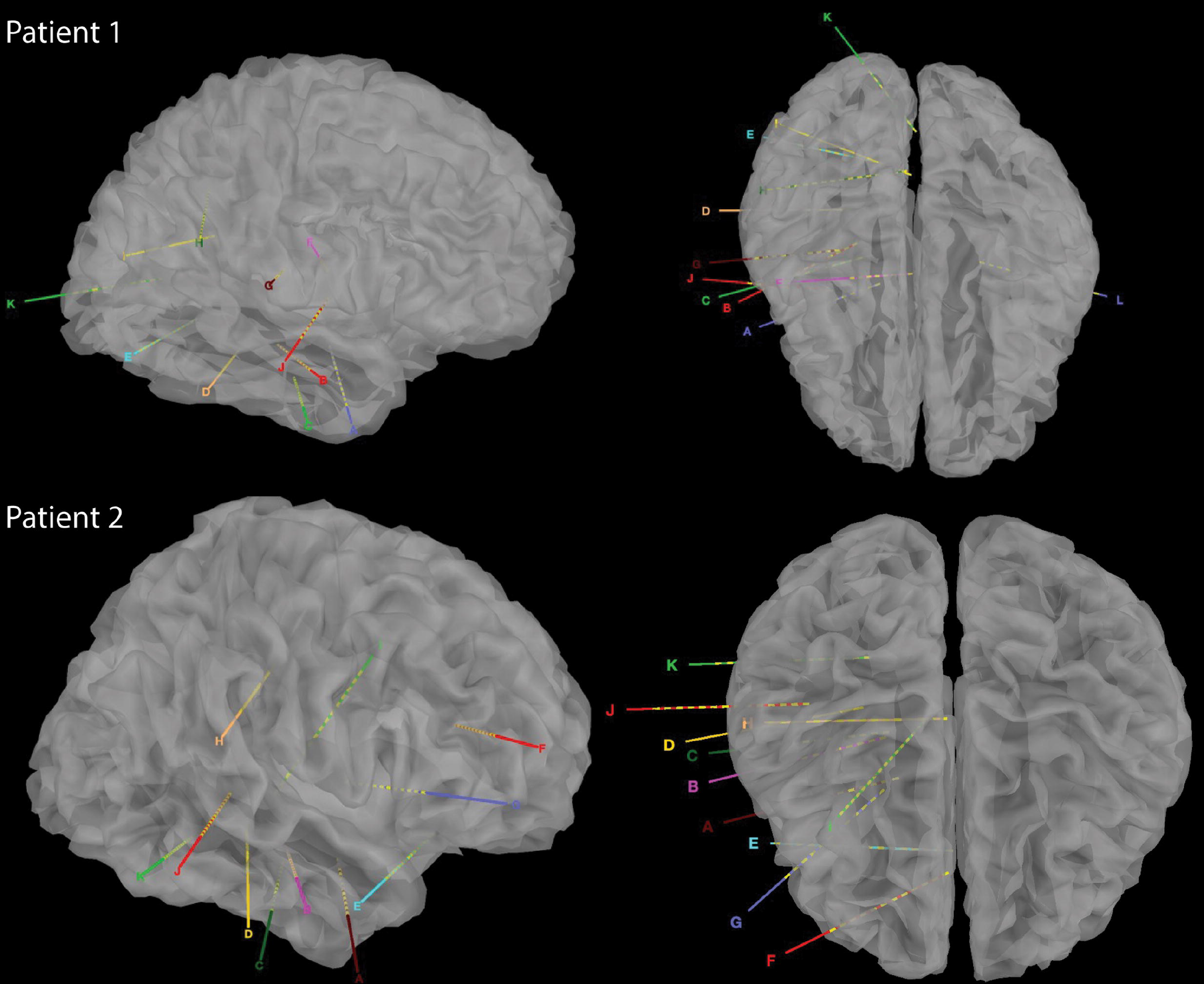
Reconstructed CT images of sEEG implants in the two patients.

During the 2-week window to localize seizure foci, we asked the patients to speak out loud when a cue was given while simultaneously recording their voice and synchronized intracranial neurophysiological signals (Figure 2). A total of 407 characters were recorded over repeated trials, covering all possible pronunciations and tones in Mandarin Chinese (Supplementary Table 1).

**Figure 2.**
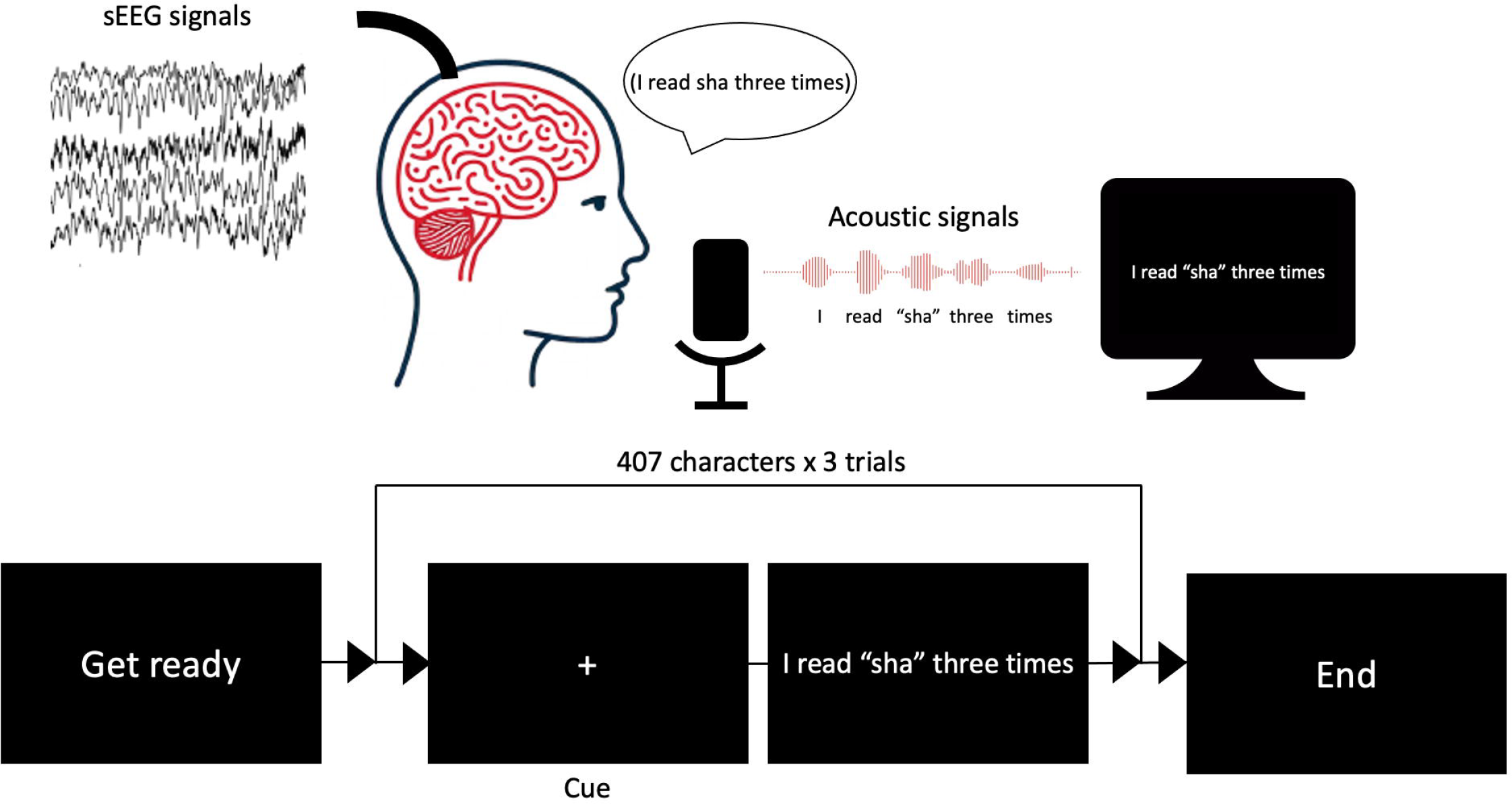
Schematic diagram of experimental design to record vocal and electrophysiological signals simultaneously.

### sEEG and acoustic signal processing

A total of 290 (148 + 142) sEEG contacts were implanted. We began signal processing by linearly detrending the sEEG signals. For extracting valuable insights from the sEEG signals, we determined the power in the 1-30 Hz, 30-70 Hz, and 70-150 Hz frequency range, which is believed to represent ensemble spiking and offers specific data about movement and speech functions. The power of the signal was ascertained by averaging the squared signal per window. We then applied transformation to the metrics to standardize their distribution. Each contact’s activity was normalized to have zero mean and a variance of one. Regarding the acoustic data, voicing of each character was semi-automatically segmented, and categorized based on vowels, consonants, and tuning. Each consonant can be assigned a corresponding set of articulatory places and manners based on international standard (Table 1)^3^. Power features of synchronized sEEG signals were segmented and categorized accordingly.

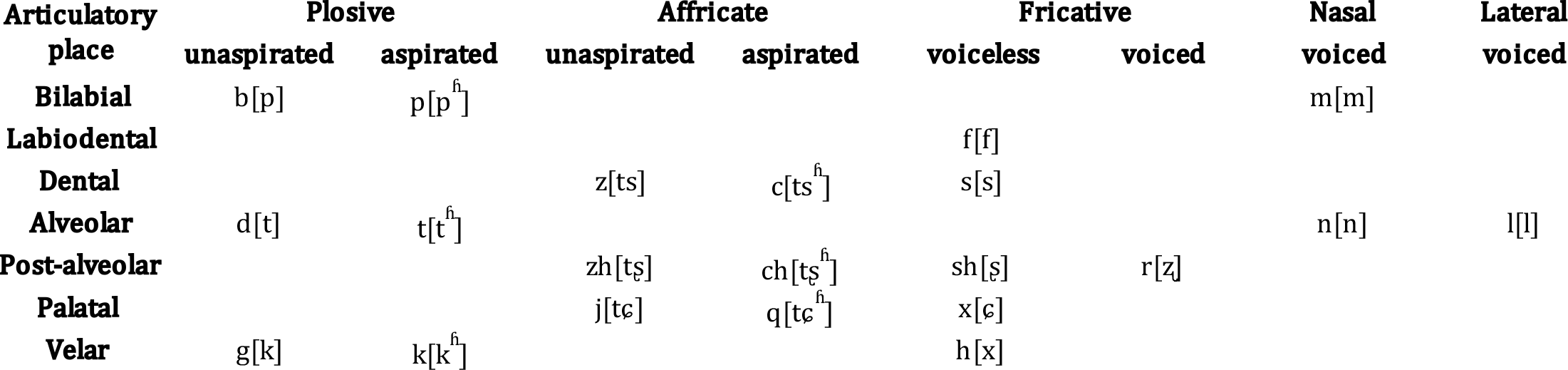

### Speech decoding from sEEG signals using recurrent neural network

We trained a long short-term memory (LSTM) recurrent neural network to interpret articulatory places, manners, and tuning. The LSTM predicted each articulatory sample using a fixed-time window of neural activity from various frequency bands, sourced from all sEEG electrodes and centered on the decoded sample. The optimization of the decoder structure involved adjusting the number of deep recurrent network layers, feedforward layers, bidirectional LSTM layers, and cells. The primary goal of training the network was to minimize the mean squared error between the decoded output and the actual output.

## RESULTS

### DECODING CONSONANTS BASED ON ARTICULATORY PLACE AND ARTICULATORY MANNER CLASSIFICATION USING SEEG SIGNALS FROM SINGLE REGION

We used sEEG 1-30 Hz, 30-70 Hz, and 70-150 Hz frequency band power of electrophysiological signals from individual brain regions to classify articulatory place and articulatory manner. The pure chance level for articulatory place and articulatory manner classification was 0.143 (1/7) and 0.125 (1/8), respectively. Our results indicated that 70-150 Hz frequency band power showed the best classification capability for both articulatory place and manner prediction across brain regions in general, which was in line with previous reports^4^. For articulatory place classification, the superior temporal gyrus showed the best performance, with an accuracy of 86.5% (Figure 3A). For articulatory manner classification, the superior temporal gyrus and the thalamus had the best results, classifying successfully 51.5% and 51.7% of the articulatory manner, respectively (Figure 3B).

**Figure 3.**
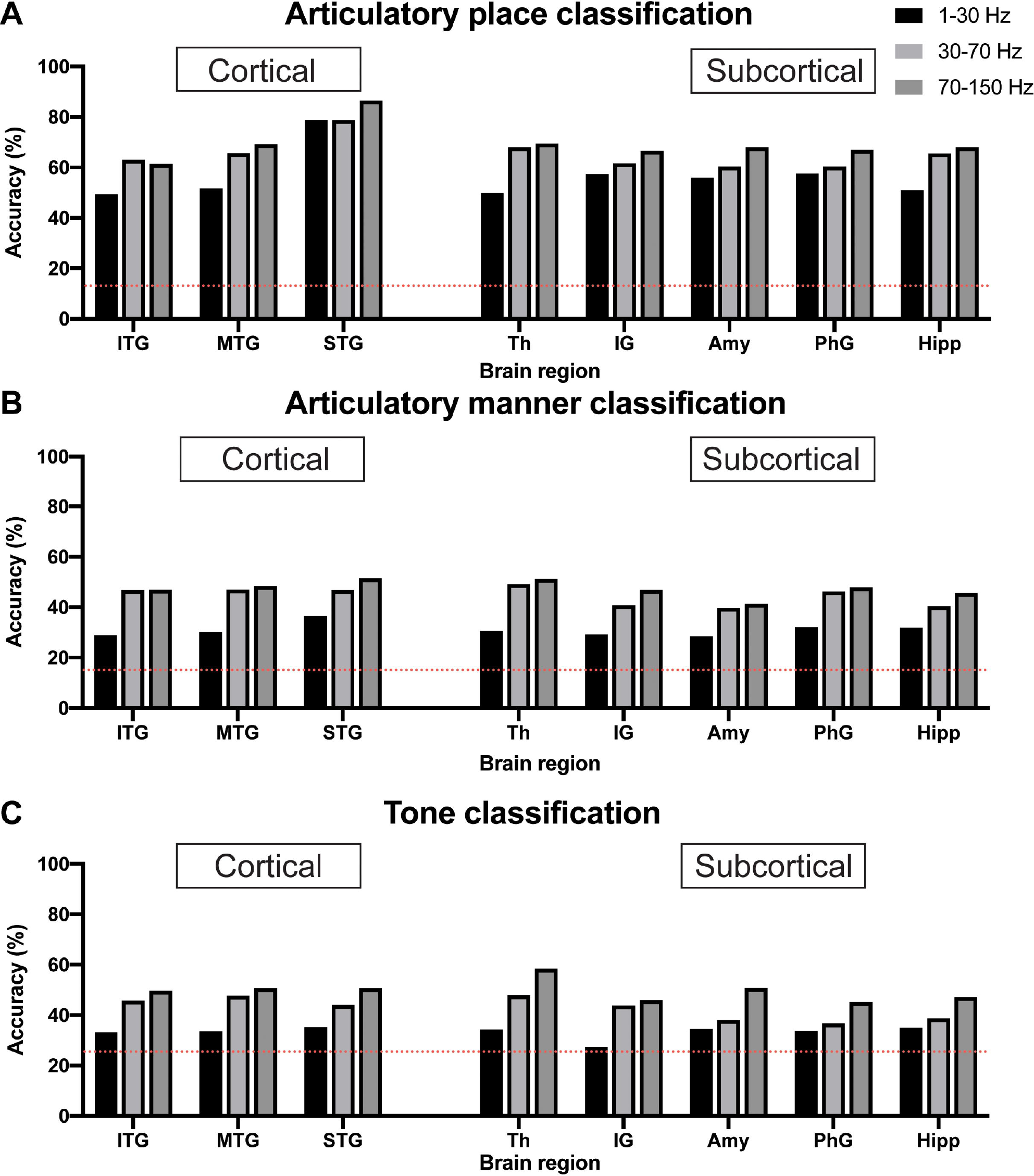
Classification accuracy for articulatory place (A), articulatory manner (B), and tone (C) using electrophysiological signals from cortical vs subcortical brain regions. Dotted red lines indicate chance levels. (A) sEEG features from the superior temporal gyrus generate the best prediction results for articulatory place. (B) sEEG features from the superior temporal gyrus and the thalamus generate the best prediction results at similar levels for articulatory manner. (C) sEEG features from the thalamus generate the best prediction results for tone. Power in the 70-150 Hz frequency band is best feature for prediction vs powers in the 1-30 Hz and 30-70 Hz frequency bands. ITP: inferior temporal gyrus, MTP: middle temporal gyrus, STP: superior temporal gyrus, Th: thalamus, IG: insular gyrus, Amy: amygdala, PhG: parahippocampal gyrus, Hipp: hippocampus.

### DECODING TONES USING SEEG SIGNALS FROM SINGLE REGION

Similar to articulatory place and manner decoding, we used 1-30 Hz, 30-70 Hz, and 70-150 Hz frequency band power of sEEG electrophysiological signals from individual brain regions to classify tone. The pure chance level for tone classification was 0.25 (1/4). Our results indicated that 70-150 Hz frequency band power still possessed the best classification capability for tone prediction across brain regions in general, and the thalamus showed the best performance, with an accuracy of 58.3% (Figure 3C).

### DECODING CONSONANTS AND TONES USING SEEG SIGNALS FROM CORTICAL AND SUBCORTICAL REGIONS COMBINED

We then used combined electrophysiological signals, one channel from cortical and one channel from subcortical brain regions, to decode consonants and tones. For articulatory place classification, we found that sEEG signals from the superior temporal gyrus were able to produce best classification results, with or without sEEG signals from subcortical regions (Figure 4A). Combining input signals from inferior temporal gyrus with hippocampus improved prediction, but still lower than what superior temporal gyrus was able to predict by itself (Figure 4B). For articulatory manner classification, sEEG signals from the superior temporal gyrus combined with signals from the thalamus were able to make best prediction (Figures 4C, D). For tone classification, sEEG signals from the thalamus profoundly improved classification results when combined with signals from the inferior, middle, and superior temporal gyri, still producing the best results when combined with the superior temporal gyrus (Figures 4E, F).

**Figure 4.**
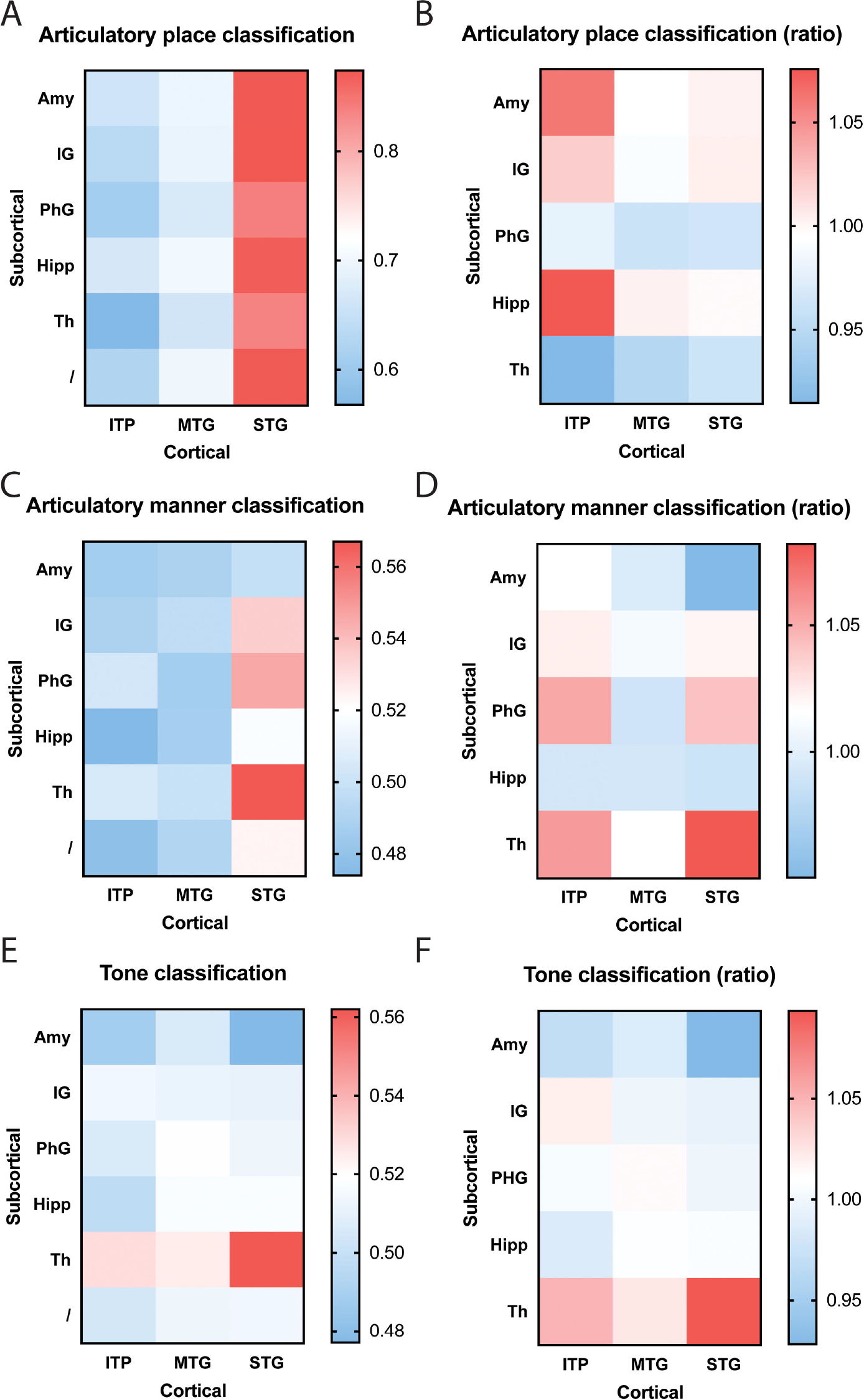
Classification accuracy and improvement ratio for articulatory place (A and B), articulatory manner (C and D), and tone (E and F) when cortical electrophysiological signals were combined with subcortical electrophysiological signals. (A and B) sEEG features from the superior temporal gyrus are best at predicting articulatory place, with or without sEEG input from subcortical regions. sEEG features from the inferior temporal gyrus may benefit from sEEG input from subcortical regions during articulatory place prediction, but its absolute accuracy remains lower than sEEG features from the superior temporal gyrus alone. (C and D) sEEG features from the superior temporal gyrus combined with sEEG features from the thalamus produce the best prediction results for articulatory manner, higher than the prediction accuracy generated from sEEG features from these two structures alone. (E and F) sEEG features from the superior temporal gyrus combined with sEEG features from the thalamus produce the best prediction results for tone, but it remains lower than the prediction accuracy generated from sEEG features from the thalamus alone. ITP: inferior temporal gyrus, MTP: middle temporal gyrus, STP: superior temporal gyrus, Th: thalamus, IG: insular gyrus, Amy: amygdala, PhG: parahippocampal gyrus, Hipp: hippocampus.

## DISCUSSION

Our work demonstrates the feasibility and value of electrophysiological signals recorded in both cortical and subcortical regions for speech decoding. Our findings are particularly significant for the design of speech neuroprostheses, as they suggest that incorporating signals from both cortical and subcortical structures could enhance the performance of these devices. The center for language processing is generally believed to be in the cortical area around the sylvian fissure of the left hemisphere called the perisylvian area. Past studies focus on harvesting signals from this area for speech decoding, while other studies have indicated the involvement of subcortical structures, such as the hippocampus and the thalamus, during speech processing^5, 6, 7, 8, 9^.

In our study, the perisylvian area, i.e. superior temporal gyrus remains highly relevant for speech decoding. We are able to use signals from the superior temporal gyrus to classify articulatory place and articulatory manner, which will help predict consonants, as well as tone classification. Signals from subcortical areas seem less relevant for articulatory place prediction, when superior temporal gyrus is used. But for articulatory area and tone predictions, signals from the thalamus substantially improve accuracy when combined with signals from the superior temporal gyrus.

Interestingly, the prediction accuracy for articulatory place is the highest, while its chance level is the lowest, compared to articulatory manner and tone. We do not have a clear explanation for this, but we believe it reflects the neural representation of the signals captured. Another interesting finding is that thalamic neural signals are best for tone prediction, which may serve as an important piece of information for research in the field of evolutionary linguistics.

Currently there are several groups investigating the use of sEEG signals for speech decoding. Angrick et al. show that sEEG and cortical-only ECoG yield similar results for speech decoding^10^. Soroush et al. study signals from grey and white matter for speech activity detection^11^. The same group also report significant contributions from deep brain structures for speech decoding^12^. Thomas et al. use sEEG approach but only include cortical regions in their study, and report neural correlates in multiple cortical regions for both articulatory and phonetic components^13^. Ramos-Escobar et al. report evidence of hippocampal involvement in the speech segmentation process^14^. Cometa et al. discovered involvement from both cortical and subcortical in syntactic processing, including from the non-dominant hemisphere^15^. Verwoert et al. published an open access sEEG dataset of 10 participants reading Dutch words^16^. Afif et al. also reported speech arrest after stimulating the insula electrically, implicating speech production in subcortical areas^17^.

Our study has limitations. The sample size, comprising only two Mandarin Chinese-speaking individuals, may limit the generalizability of our findings. Additionally, the study’s focus on right hemisphere regions could miss critical information processed in the left hemisphere, traditionally associated with language. Furthermore, the clinical condition of our participants (refractory epilepsy) and the resulting altered neurophysiology could affect the generalizability of our findings to the broader population.

Looking forward, our research opens several avenues for further investigation. Larger-scale studies involving diverse languages and larger participant cohorts could validate and extend our findings. Moreover, longitudinal studies could examine the stability of sEEG signal decoding over time, which is crucial for the practical application of BCIs in chronic conditions. Finally, integrating our findings with machine learning advancements could lead to more sophisticated and accurate speech neuroprosthesis designs, ultimately enhancing the quality of life for individuals with speech impairments.

In conclusion, our study represents a significant step towards understanding and harnessing the full potential of brain signals for speech decoding. The implications for assistive technologies are profound, offering a chance for restoring communication abilities to those who have lost them.

## Data availability statement

The raw data supporting the conclusions of this article will be made available by the authors, without undue reservation.

## Ethics statement

This study was approved by the Ethics Committee of Zhejiang University School of Medicine Second Affiliated Hospital (I2022145). It was conducted in accordance with the local legislation and institutional requirements.

## Author contributions

HW was responsible for the writing of this paper. CC, WM, and CF were responsible for the experimental collection of original data. WC, ZZhu, HJ, ZZheng, and MS were responsible for the data collection and analysis. HW, TW, and JZ were responsible for the experimental design and overall planning of the research. All authors contributed to the article and approved the submitted version.

## Acknowledgments

We would like to thank the participants who believed in us and volunteered in this study.

