## Supplementary Table 1 for "Investigation of contributions from cortical and subcortical brain structures for speech decoding"

| | a | o | e | i | -i | u | ü | ai | ei | ui | ao | ou | iu | ie | üe | er | an | en | in | un | ün | ang | eng | ing | ong | i-a | i-an | i-ang | i-ao | i-ong | u-a | u-ai | u-an | u-ang | u-o | ü-an |
|---|---|---|---|---|---|---|---|---|---|---|---|---|---|---|---|---|---|---|---|---|---|---|---|---|---|---|---|---|---|---|---|---|---|---|---|---|
| b | ba | bo | | bi | | bu | | bai | bei | | bao | | | bie | | | ban | ben | bin | | | bang | beng | bing | | | bian | | biao | | | | | | | |
| p | pa | po | | pi | | pu | | pai | pei | | pao | pou | | pie | | | pan | pen | pin | | | pang | peng | ping | | | pian | | piao | | | | | | | |
| m | ma | mo | me | mi | | mu | | mai | mei | | mao | mou | miu | mie | | | man | men | min | | | mang | meng | ming | | | mian | | miao | | | | | | | |
| f | fa | fo | | | | fu | | | fei | | | fou | | | | | fan | fen | | | | fang | feng | | | | | | | | | | | | | |
| d | da | | de | di | | du | | dai | dei | dui | dao | dou | diu | die | | | dan | den | | dun | | dang | deng | ding | dong | dia | dian | | diao | | | | duan | | duo | |
| t | ta | | te | ti | | tu | | tai | | tui | tao | tou | | tie | | | tan | | | tun | | tang | teng | ting | tong | | tian | | tiao | | | | tuan | | tuo | |
| n | na | | ne | ni | | nu | nü | nai | nei | | nao | nou | niu | nie | nüe | | nan | nen | nin | | | nang | neng | ning | nong | | nian | niang | niao | | | | nuan | | nuo | |
| l | la | lo | le | li | | lu | lü | lai | lei | | lao | lou | liu | lie | lüe | | lan | | lin | lun | | lang | leng | ling | long | lia | lian | liang | liao | | | | luan | | luo | |
| g | ga | | ge | | | gu | | gai | gei | gui | gao | gou | | | | | gan | gen | | gun | | gang | geng | | gong | | | | | | gua | guai | guan | guang | guo | |
| k | ka | | ka | | | ku | | kai | kei | kui | kao | kou | | | | | kan | ken | | kun | | kang | keng | | kong | | | | | | kua | kuai | kuan | kuang | kuo | |
| h | ha | | he | | | hu | | hai | hei | hui | hao | hou | | | | | han | hen | | hun | | hang | heng | | hong | | | | | | hua | huai | huan | huang | huo | |
| j | | | | ji | | | ju | | | | | | jiu | jie | jue | | | | jin | | jun | | | jing | | jia | jian | jiang | jiao | jiong | | | | | | juan |
| q | | | | qi | | | qu | | | | | | qiu | qie | que | | | | qin | | qun | | | qing | | qia | qian | qiang | qiao | qiong | | | | | | quan |
| x | | | | xi | | | xu | | | | | | xiu | xie | xue | | | | xin | | xun | | | xing | | xia | xian | xiang | xiao | xiong | | | | | | xuan |
| zh | zha | | zhe | | (zhi) | zhu | | zhai | zhei | zhui | zhao | zhou | | | | | zhan | zhen | | zhun | | zhang | zheng | | zhong | | | | | | zhua | zhuai | zhuan | zhuang | zhuo | |
| ch | cha | | che | | (chi) | chu | | chai | | chui | chao | chou | | | | | chan | chen | | chun | | chang | cheng | | chong | | | | | | chua | chuai | chuan | chuang | chuo | |
| sh | sha | | she | | (shi) | shu | | shai | shei | shui | shao | shou | | | | | shan | shen | | shun | | shang | sheng | | | | | | | | shua | shuai | shuan | shuang | shuo | |
| r | | | re | | (ri) | ru | | | | rui | rao | rou | | | | | ran | ren | | run | | rang | reng | | rong | | | | | | | | ruan | | ruo | |
| z | za | | ze | | (zi) | zu | | zai | zei | zui | zao | zou | | | | | zan | zen | | zun | | zang | zeng | | zong | | | | | | | | zuan | | zuo | |
| c | ca | | ce | | (ci) | cu | | cai | | cui | cao | cou | | | | | can | cen | | cun | | cang | ceng | | cong | | | | | | | | cuan | | cuo | |
| s | sa | | se | | (si) | su | | sai | | sui | sao | sou | | | | | san | sen | | sun | | sang | seng | | song | | | | | | | | suan | | suo | |
| y | ya | yo | | | (yi) | | (yu) | | | | yao | you | | (ye) | (yue) | | yan | | (yin) | | (yun) | yang | | (ying) | yong | | | | | | | | | | | (yuan) |
| w | wa | wo | | | | (wu) | | wai | wei | | | | | | | | wan | wen | | | | wang | weng | | | | | | | | | | | | | |
